# Neonatal enthesis healing involves non-inflammatory formation of acellular scar through ECM secretion by resident cells

**DOI:** 10.1101/2021.12.20.473454

**Authors:** Ron Carmel Vinestock, Neta Felsenthal, Eran Assaraf, Eldad Katz, Sarah Rubin, Lia Heinemann-Yerushalmi, Sharon Krief, Nili Dezorella, Smadar Levin-Zaidman, Michael Tsoory, Stavros Thomopoulos, Elazar Zelzer

## Abstract

Wound healing is a well-orchestrated process that typically recruits the immune and vascular systems to restore the structure and function of the injured tissue. Injuries to the enthesis, a hypocellular and avascular tissue, often result in fibrotic scar formation and loss of mechanical properties, thereby severely affecting musculoskeletal function and life quality. This raises questions about the healing capabilities of the enthesis.

Here, we established an injury model to the Achilles entheses of neonatal mice to study the possibility that at an early age, the enthesis can heal more effectively. Histology and immunohistochemistry analyses revealed an atypical process that did not involve inflammation or angiogenesis. Instead, neonatal enthesis healing was mediated by secretion of collagen types I and II by resident cells, which formed a permanent hypocellular and avascular scar. Transmission electron microscopy showed that the cellular response to injury, including ER stress, autophagy and cell death, varied between the tendon and cartilage ends of the enthesis. Single-molecule *in situ* hybridization, immunostaining, and TUNEL assays verified these differences. Finally, gait analysis showed that these processes effectively restored function of the injured leg.

Collectively, these findings reveal a novel healing mechanism in neonatal entheses, whereby local ECM secretion by resident cells forms an acellular ECM deposit in the absence of inflammation markers, allowing gait restoration. These insights into the healing mechanism of a complex transitional tissue may lead to new therapeutic strategies for adult enthesis injuries.

## INTRODUCTION

Wound healing is a critical and complex process that restores structure and function by replacing damaged tissue. In adult animals, this process comprises a sequential cascade of overlapping events, including bleeding and activation of the coagulation system, recruitment of inflammatory cells, fibroblast migration, collagen synthesis, angiogenesis, and remodeling of the injury site^1,2^. In the musculoskeletal system, however, tissues differ in their ability to heal injuries^3^. Whereas bones and muscles have regenerative capacities^4,5^, tendon, ligament, and cartilage tissues often heal via scar formation, without complete restoration of mechanical properties. Interestingly, these scar-forming tissues are all extracellular matrix (ECM)-rich, hypocellular, poorly vascularized, and slow proliferating, all of which may influence the repair process^6–8^.

Tendon, ligament, and cartilage healing have been investigated using a variety of injury models in different organisms^6,9^. In tendons, common injury models include full or partial transection of the tendon and overuse injuries caused by physical activities^10–12^. During the healing process, recruited fibroblasts initially synthesize predominantly collagen type III, later replacing it with collagen type I. However, this remodeling process is typically insufficient, often resulting in an altered ECM composition compared to the uninjured tissue^13^. Thus, the repair of injured adult tendons usually involves formation of a fibrovascular scar and loss of histological and mechanical characteristics^13–15^. However, a recent study in neonatal mice showed regenerative properties of Achilles tendon after transection^10^.

Studies of cartilage repair have established different healing responses in partial-thickness compared to full-thickness injuries. Partial-thickness injuries do not heal spontaneously, as inflammation and blood vessels are absent^16^. Following injury, cells within the wound margins undergo chondroptosis, a variant of cell death that is characterized by an increase in Golgi apparatus and endoplasmic reticulum (ER), autophagic vacuoles, patchy condensations of nuclei and blebbing of cytoplasmic material and activation of apoptosis via caspase-3 and caspase-9 involvement^17–22^. In full-thickness injuries, the damage is deeper and reaches into the subchondral bone, thereby creating a pathway into the vascular bone marrow^23^. This enables access for the recruitment of immune cells, eventually leading to fibrocartilage formation and ECM deposition in the wound area^24^. As the newly formed fibrocartilage is weaker than the original tissue, the cartilage undergoes gradual degradation, which leads to loss of mechanical properties^23–25^.

Another component of the musculoskeletal system is the enthesis, a fibrocartilaginous tissue that bridges tendon and bone. This unique specialized connective tissue attaches the two distinct tissues by forming a gradient of cellular and extracellular features along its length^26,27^. Adult enthesis repair commonly results in permanent damage due to failure to restore its structure. The resulting mechanically insufficient attachment limits joint function and is prone to retears^28^. However, the outcome may depend on the severity and type of injury. Comparison of the mechanical outcomes of rotator cuff injuries in adult mice suggested that repair is more successful after partial injuries, including reduced scar formation and recovery of gait^29^. Furthermore, it was suggested that rotator cuff entheses of neonatal mice display some regenerative capacity^30^. The different healing potential between adult and neonatal mice opens the possibility that the healing capacity of the enthesis is switched off early after birth. Moreover, given the complexity of the enthesis structure, it is unclear whether the healing process, if exists, follows the same course in all enthesis zones.

To address these questions, we turned to the Achilles enthesis of neonatal mouse as a model and we induced a needle punch injury to this enthesis. Notably, we took great care not to punch all the way through the enthesis and into the bone marrow space. Although we observed no inflammation or angiogenesis at the injury site, temporal analyses identified the formation of an acellular domain and ECM plug at the injury site. The acellular domain was composed of collagen type I at the tendon end and collagen type II at the cartilage end, suggesting that this domain is formed locally by the resident enthesis cells. Immunostaining, gene expression analyses, TUNEL assay, and transmission electron microscopy revealed that cells at the injury site undergo ER stress and autophagy, and suggested that the observed cellular loss was a result of chondroptosis-like cell death. Gait recovery suggested that, despite the loss of tissue structure, the healing process effectively restored joint function. Together, these findings reveal in neonatal mice a novel mechanism whereby extensive secretion of ECM by resident cells at the injury site drives enthesis healing and restores its function.

## RESULTS

### The neonatal enthesis heals by formation of hypocellular domains flanking an ECM plug

To study the healing process of the enthesis, we established a mouse model of partial injury (Fig. 1A). Using postnatal day (P) 7 mice, we performed a minimal cut of the skin and inserted a thin 32G needle through the tendon and the Achilles enthesis, reaching the distal part of the calcaneus. This procedure injured the enthesis throughout its length, while avoiding penetration of the bone marrow cavity and bleeding, thereby excluding the potential contribution of cells from the marrow space. Notably, as the mouse enthesis mineralizes at approximately two weeks postnatally, the injury was made in an unmineralized enthesis.

**Figure 1.**
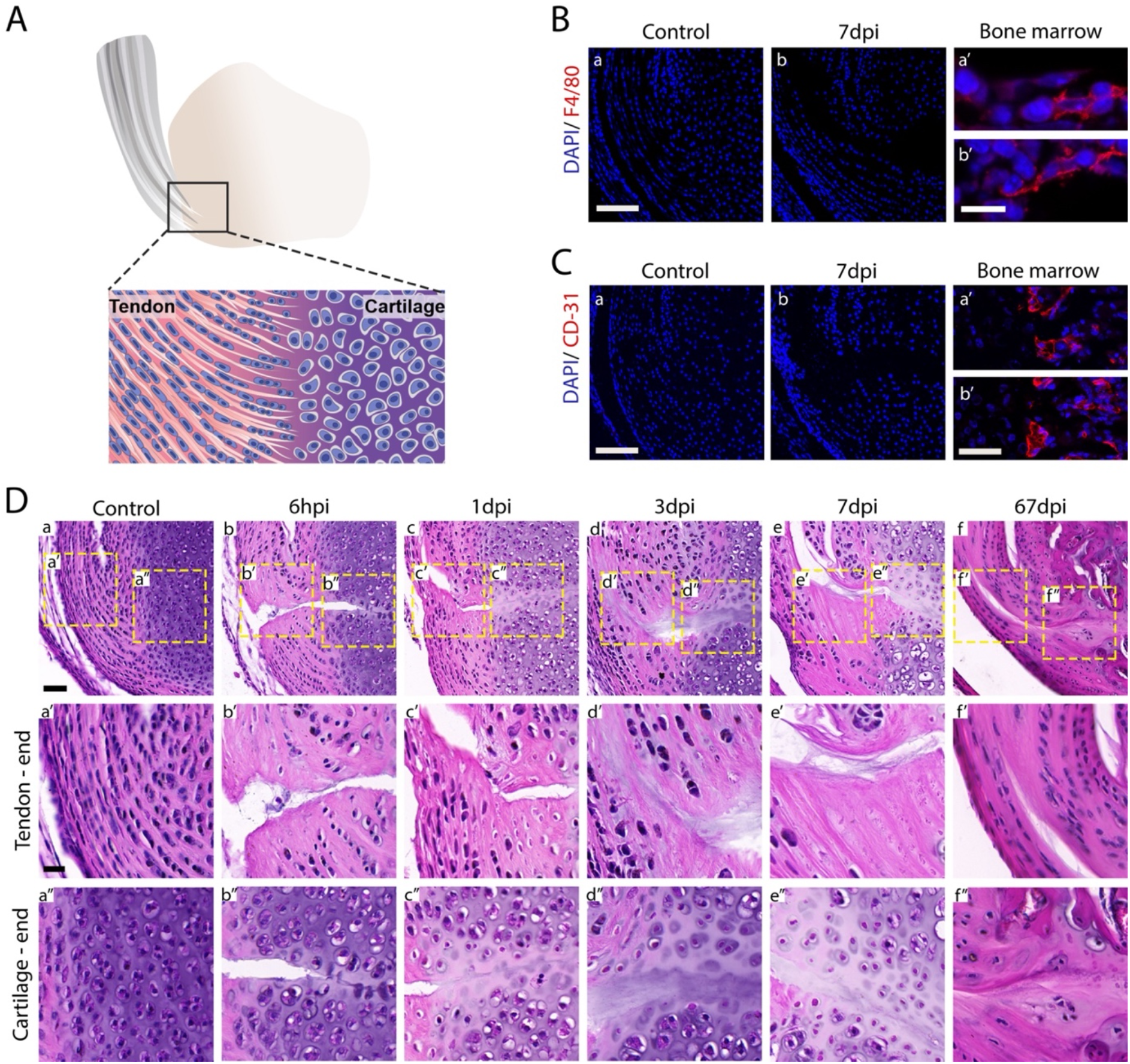
Enthesis healing involves formation of hypocellular domains flanking an ECM plug without typical features of wound healing. **A** Schematic illustration of the Achilles enthesis. **B** F4/80 immunohistochemistry staining of sections through the enthesis at 7 dpi shows no macrophages in proximity to the injury site (a,b). **C** CD31 immunohistochemistry staining at 7 dpi shows no angiogenesis at the injury site (a,b). Bone marrow was used as a control (a’, uninjured leg; b’, injured leg). **D** Hematoxylin and eosin staining of sagittal sections of P7 control uninjured entheses (a) and injured entheses at 6 hours (hpi) to 67 days (dpi) post-injury (b-f). Scale bar: 50 μm. Ca’-f and Ca”-f” are magnifications of the tendon end and cartilage end areas in Ca-f, respectively. Scale bar: 20 μm.

Following the classical model of wound healing, we examined the injury site for the presence of inflammation. For that, day 7 post-injury and non-injured limb sections were immunostained for F4/80, a specific marker for macrophages^31^. As seen in Figure 1B, no macrophages were detected in or around the injury site. To determine whether there was angiogenesis in the injured enthesis, we stained for the endothelial marker CD31^32^ (Fig. 1C). The results showed no blood vessel invasion into the injury site. Together, these results suggested that in our enthesis injury model, healing did not involve inflammation and angiogenesis.

To uncover the temporal profile of the healing process in the injured enthesis, we analyzed histological sections of tissues at 6 hours, 1-, 3-, 7- and 67-days post-injury (dpi). At 6 hours, the injury site was clearly observed as a cut that extended between the Achilles tendon across the enthesis to the distal cartilage end of the calcaneus (Fig. 1Db-b”). Interestingly, at day 3 post-injury, in the region of the injury site near the tendon, an acellular domain was forming and propagating several cell rows away from the injury margins. At the yet unmineralized cartilage end of the enthesis, a similar propagation of the response area away from the injury site was observed. Chondrocytes adjacent to the injury site lost most of their volume, while the matrix around them appeared abnormal, as it was paler than in the control. Within the injury site, there was an accumulation of ECM that filled part of the gap, forming a plug-like structure (Fig. 1Dd-d”). In line with the macrophage and endothelial cell results (Fig. 1B,C), there were no signs of inflammation or angiogenesis at the injury site. At 7 dpi, the injury site was still recognizable, with the area around it maintaining its acellular characteristics and an ECM plug filling the gap (Fig. 1De-e”). This morphology of the injury site was unchanged at 67 dpi (Fig. 1Df-f”).

To evaluate long-term remodeling of the injured enthesis, we examined mineralization by staining sections with Von Kossa stain at 5 months post-injury. As seen in Figure S1, the healed enthesis was mineralized; however, the tidemark, which is generally considered the border between mineralized and non-mineralized parts of the fibrocartilage enthesis, was disrupted, indicating a failure of the enthesis to heal completely.

Collectively, the absence of inflammation and angiogenesis and the formation of hypocellular domains flanking an ECM plug suggest that the enthesis can heal via a novel mode of tissue repair.

### Hypocellular scar at the healing neonatal enthesis is formed locally by resident cells

Previous studies on enthesis healing described the formation of a scar tissue enriched with collagen type III^33,34^. To examine if the observed hypocellular scar in the injury site was composed of collagen type III, we immunostained 3 dpi and control limbs for COL3A1. As seen in Figure S2, we failed to observe changes in the level of COL3A1 in or around the injury site.

The two main components of the enthesis ECM are collagen type I (COL1A1), which is expressed in the tendon and its transition into fibrocartilage, and collagen type II (COL2A1), which is expressed in the fibrocartilage and its transition into bone. Having observed the formation of an ECM plug in the healing enthesis, we examined the distribution of these two collagens during the healing process. For that, double-immunofluorescence staining for COL1A1 and COL2A1 was performed on sections of 3-dpi and age-matched uninjured Achilles entheses. In control entheses, the expected patterns of expression were seen, as COL1A1 was expressed in the tendon and the enthesis whereas COL2A1 was expressed across cartilage, with limited overlap between the domains (Fig. 2a). In the injured enthesis, the hypocellular domain and ECM plug were positive for COL1A1 and COL2A1 and the overall patterns of expression were maintained, suggesting that the ECM plug forms locally by the cells flanking the injury site. Nevertheless, we observed several differences between the injured and uninjured enthesis. In the former, COL2A1 expression expanded into the tendon end of the enthesis (Fig. 2 Ca-a’,b-b’), whereas in the ECM plug, the two expression domains expanded into each other.

**Figure 2.**
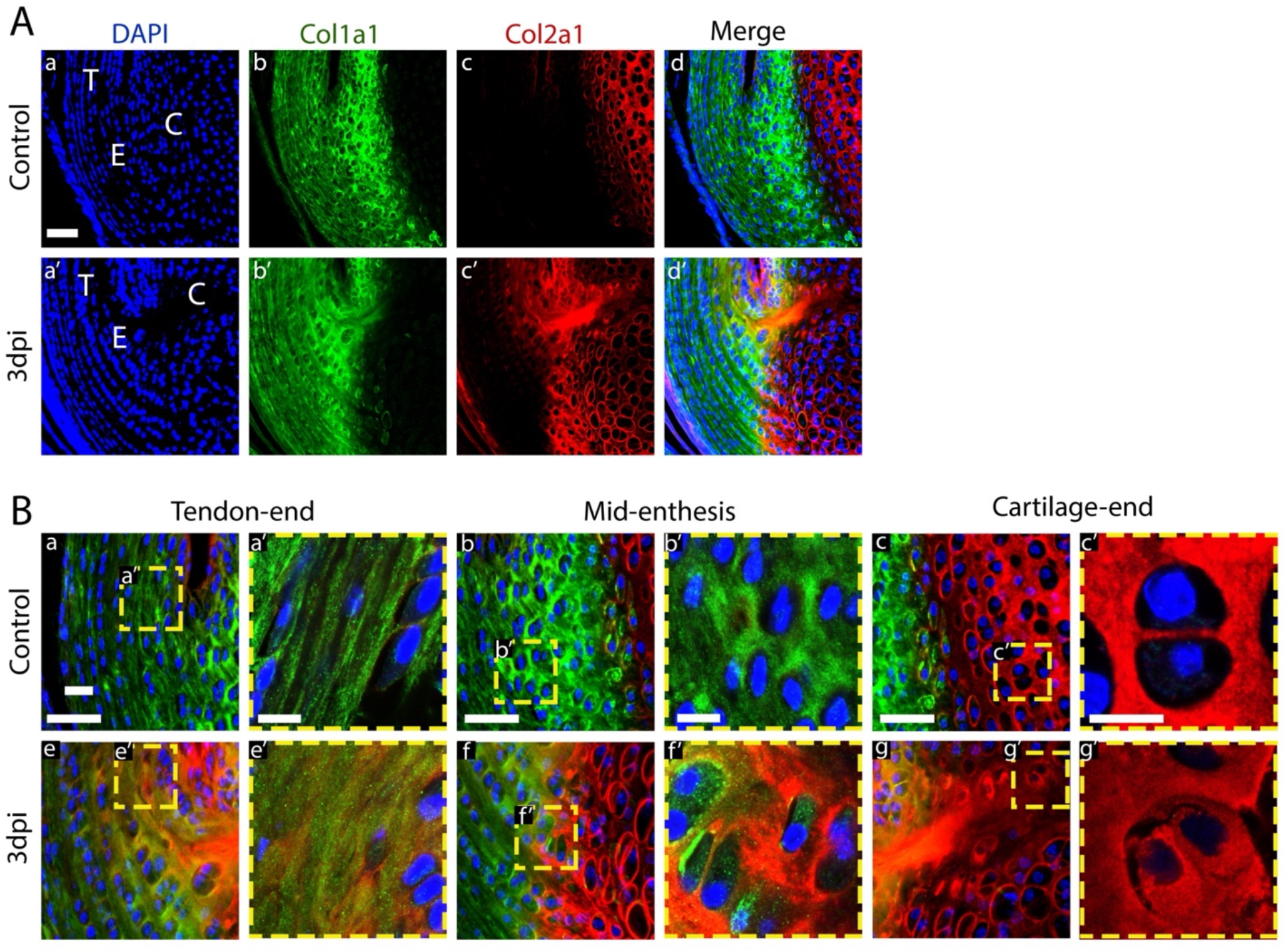
Enthesis cells form a local acellular scar by secreting COL1A1 and COL2A1. **A** Immunohistochemistry staining for COL1A1 and COL2A1 shows ECM plug and loss of expression domain in 3 dpi enthesis (a’-d’) in comparison to the control (a-d). Scale bar: 50 μm. **B** Magnifications of tendon, enthesis and cartilage in control (a-c) and 3 dpi (e-g) entheses. Scale bar: 20 μm. a’-g’ are magnifications of the boxed areas in a-g, respectively. Scale bar: 10 μm.

To gain a better understanding of the loss of segregation between COL1A1 and COL2A1 expression domains, we examined at higher magnification the cells at the border between the two domains. As seen in Figure 2B c-c’,d-d’, whereas cells in the uninjured enthesis expressed only one type of collagen, we observed same-stage cells of injured enthesis that co-expressed COL1A1 and COL2A1. In addition, whereas in the uninjured enthesis COL2A1 staining was observed in the ECM around the cells, at the cartilage end adjacent to the injury site, where histologically we identified shrinking chondrocytes, the empty lacunae of these cells were mostly COL2A1-positive and DAPI-negative.

These results suggest that the hypocellular domain around the injury site is formed by ECM secretion by neighboring local cells, which divides this domain into two regions containing either tendon or cartilage ECM. At the border between them, cells co-express COL1A1 and COL2A1, resulting in mixed ECM. While the composition of the ECM plug seems to follow the same pattern, the border between expression domains was less defined.

### Transmission electron microscopy analysis showed swollen ER, autophagic bodies and cell death along with disorganized ECM in the healing enthesis

Our finding that the healing process of the enthesis involves local ECM secretion and cell loss led us to study this process at subcellular resolution. To this end, we used transmission electron microscopy (TEM) to image both ends of the enthesis at 1- and 3-days post-injury (Fig. 3). At the tendon end of an uninjured control enthesis, we observed typical elongated tenocytes with apparent nuclei and ER (Fig. 3 Aa-a’). In contrast, 1-dpi cells of injured entheses displayed an increased number of Golgi apparatus, swollen ER, and numerous vesicles with the morphology of autophagic bodies (Fig. 3 Ab-b’). At 3 dpi, the condition of these cells deteriorated, as lost nuclei, swollen ER, and damaged mitochondria were observed (Fig. 3 Ac-c’).

**Figure 3.**
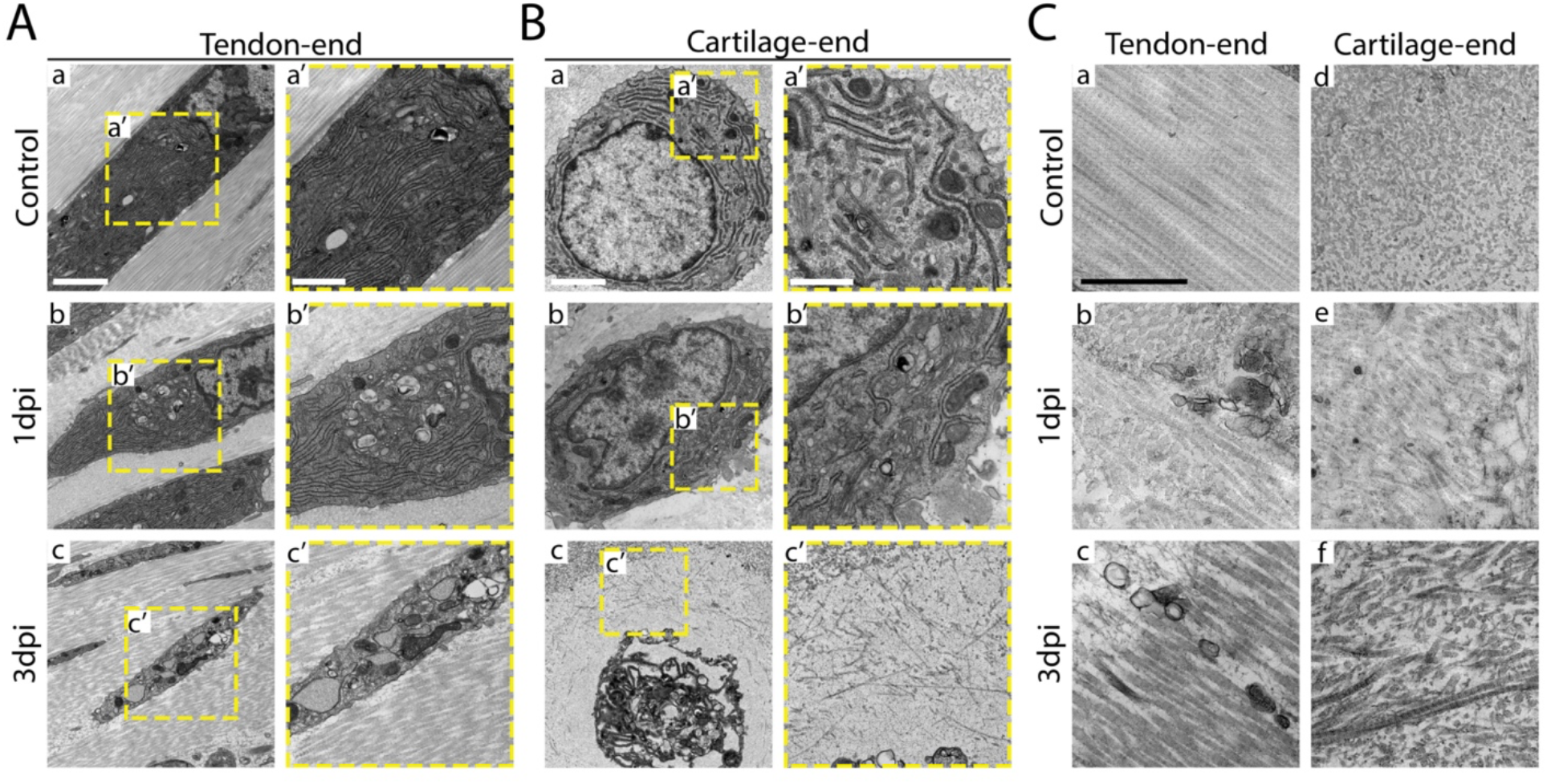
Transmission electron microscopy reveals impaired cellular and ECM morphology in the healing enthesis. **A** Cellular morphology at the tendon end of control, 1-dpi and 3-dpi entheses. (a-c) Scale bar: 2 μm. a’-c’ are magnifications of the boxed areas in a-c, respectively. Scale bar: 1 μm. **B** Cellular morphology at the cartilage end of control, 1-dpi and 3-dpi entheses. (a-c) Scale bar: 2 μm. a’-c’ are magnifications of the boxed areas in a-c, respectively. Scale bar: 1 μm. **C** Fiber orientation of the ECM plug at the tendon end (a-c) and cartilage end (d-f) of control, 1-dpi and 3-dpi entheses. Scale bar: 1 μm.

At the cartilage end of control enthesis, cells had a typical round shape of chondrocytes, with clear nuclei and extensive and well-organized ER (Fig. 3 Ba-a’). However, adjacent to the injury site at 1 dpi, cells started to detach from the ECM around them and their membranes were ruptured, causing spillage of cell contents and membrane shedding. Internally, we observed less ER, nuclei became condensed, and mitochondria appeared damaged compared to control cells. As in the cells at the tendon end, we observed autophagic bodies, swollen ER, and an increase in the number of Golgi apparatus; yet, the severity of these cellular pathologies was reduced relative to the opposite side (Fig. 3 Bb-b’). By 3 dpi, the condition of these cells dramatically deteriorated, as they lost their membranes as well as most of their organelles, including nuclei. The remains of the cell bodies were condensed in the center of their ECM lacunae. Interestingly, in the space that was left by the shrinking cells, ECM deposition was observed. This fits well with our finding of COL2A1 staining in the cell-free domains (Fig. 3 Bc-c’).

We next focused on the newly formed hypocellular domain and the ECM plug. At the tendon end of 1-dpi injured enthesis, unlike in the control, the collagen fibers were oriented in different directions (Figure 3C). At 3 dpi, collagen fibers remained disorganized, yet some alignment was observed. At the cartilage end, the ECM in the control enthesis was characterized by well-organized fibers. In contrast, in both 1-dpi and 3-dpi injured entheses, the collagen fibers were misaligned, disorganized, and unevenly distributed.

Overall, the findings of increased Golgi apparatus, swollen ER, and autophagic bodies in the injured enthesis cells are consistent with extensive ECM production, which may also explain ECM disorganization as well as induction of ER stress and autophagy. The observation of shrinking cells that had lost their nuclei suggests that these cells underwent apoptotic cell death. Finally, these results indicate that the healing process involved different responses at the two ends of the enthesis.

### The injured enthesis undergoes ER stress at 1 day post-injury

An increased secretory load during the formation of the ECM plug is expected to induce ER stress, as suggested by our finding of swollen ER in cells near the injury site. To address this possibility, we measured the expression of ER stress markers *BiP* and *CHOP*. For that, mRNA was isolated from 1-dpi Achilles entheses and uninjured controls. qRT-PCR revealed a significant elevation of both markers in the injured tissues compared to the control, suggesting that cells at the injured enthesis undergo ER stress. To verify this result and to provide spatial information about the cells that undergo ER stress, we performed single-molecule *in situ* hybridization (smFISH) for both markers on sections of 1 dpi and control entheses (Fig. 4B). Quantification showed that both markers were significantly elevated at the tendon end of the enthesis, whereas at the cartilage end only *BiP* was significantly increased (Fig. 4C).

**Figure 4.**
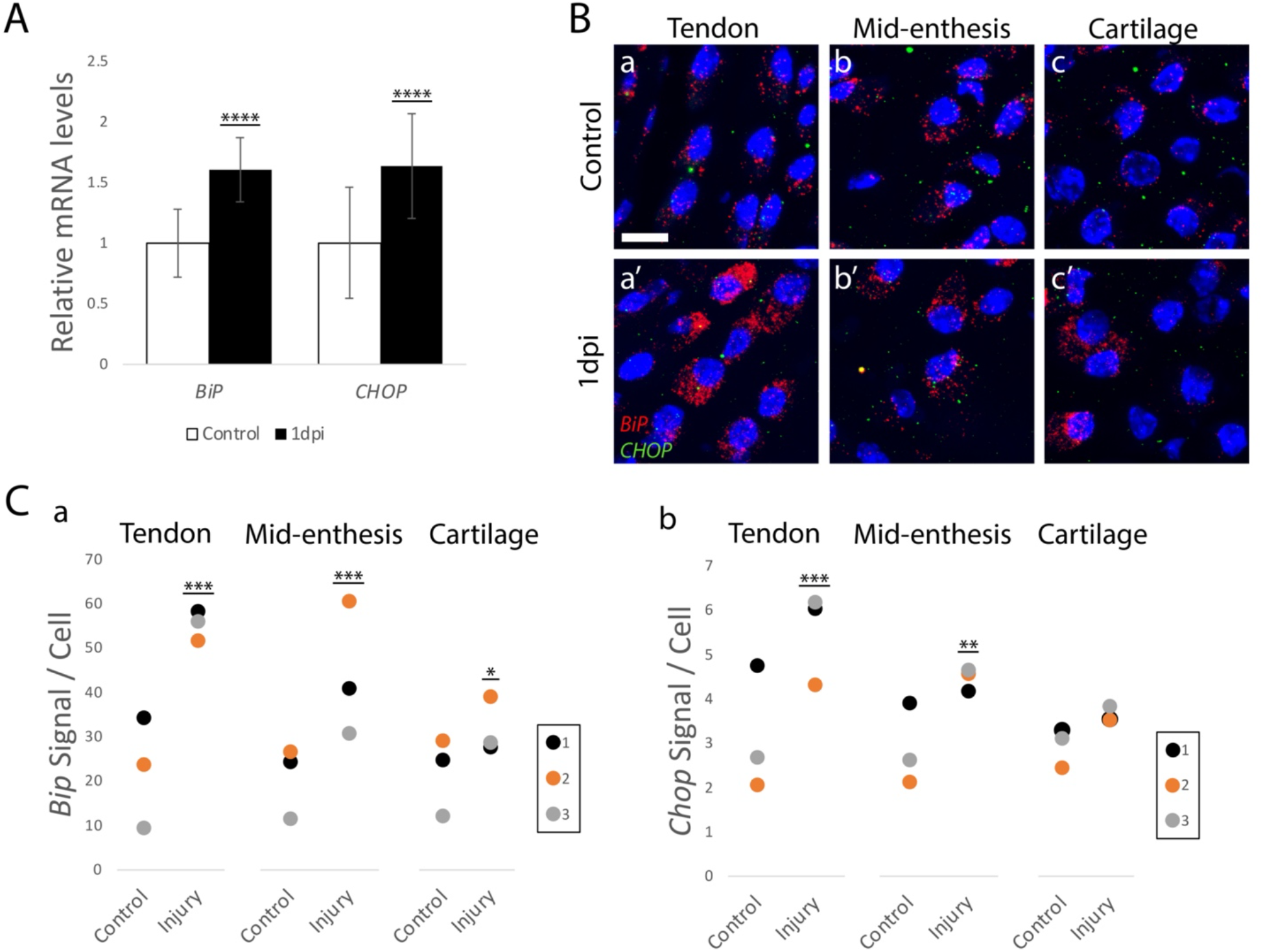
Upregulation in ER stress markers in 1-day post-injury enthesis. **A** Graph showing qRT-PCR analysis for *BiP* and *CHOP* mRNA in control and 1-dpi entheses (****, p<0.0001; n=7; data are normalized to *18S* and presented as mean±SD). **B** smFISH for ER stress markers *BiP* and *CHOP* reveals an elevation in expression levels in 1-dpi enthesis. Scale bar: 10 μm **C** Quantification of *in situ* HCR signal of *BiP* (a) and *CHOP* (b) to cell in tendon, enthesis and cartilage regions of control and 1-dpi animals (***, p<0.001; **, p<0.01; *, p<0.05; n=3; data are presented as mean). Color coding indicates paired comparison samples.

These results confirmed the induction of ER stress in the enthesis during the healing process, supporting the notion that enthesis cells were under a heavy burden of ECM secretion. Further, the results highlighted key difference between the healing responses on the two sides of the enthesis.

### Enthesis healing involves autophagy and cell death

Our TEM analysis identified cellular features that are consisted with autophagy and cell death in the injured tissue. To confirm the occurrence of these processes, we tracked molecular markers for these processes. To examine the expression of the autophagy marker LC3-II, we applied our injury procedure to GFP-LC3 mice^35^ and analyzed their entheses at 1 dpi. As seen in Figure 5A, many LC3-II positive cells were observed at the tendon end, whereas few LC3-II positive cells were observed in the middle of the enthesis and cartilage. These results support the TEM observation and suggest that the healing process involves autophagy, most prominently at the tendon end of the injured tissue.

**Figure 5.**
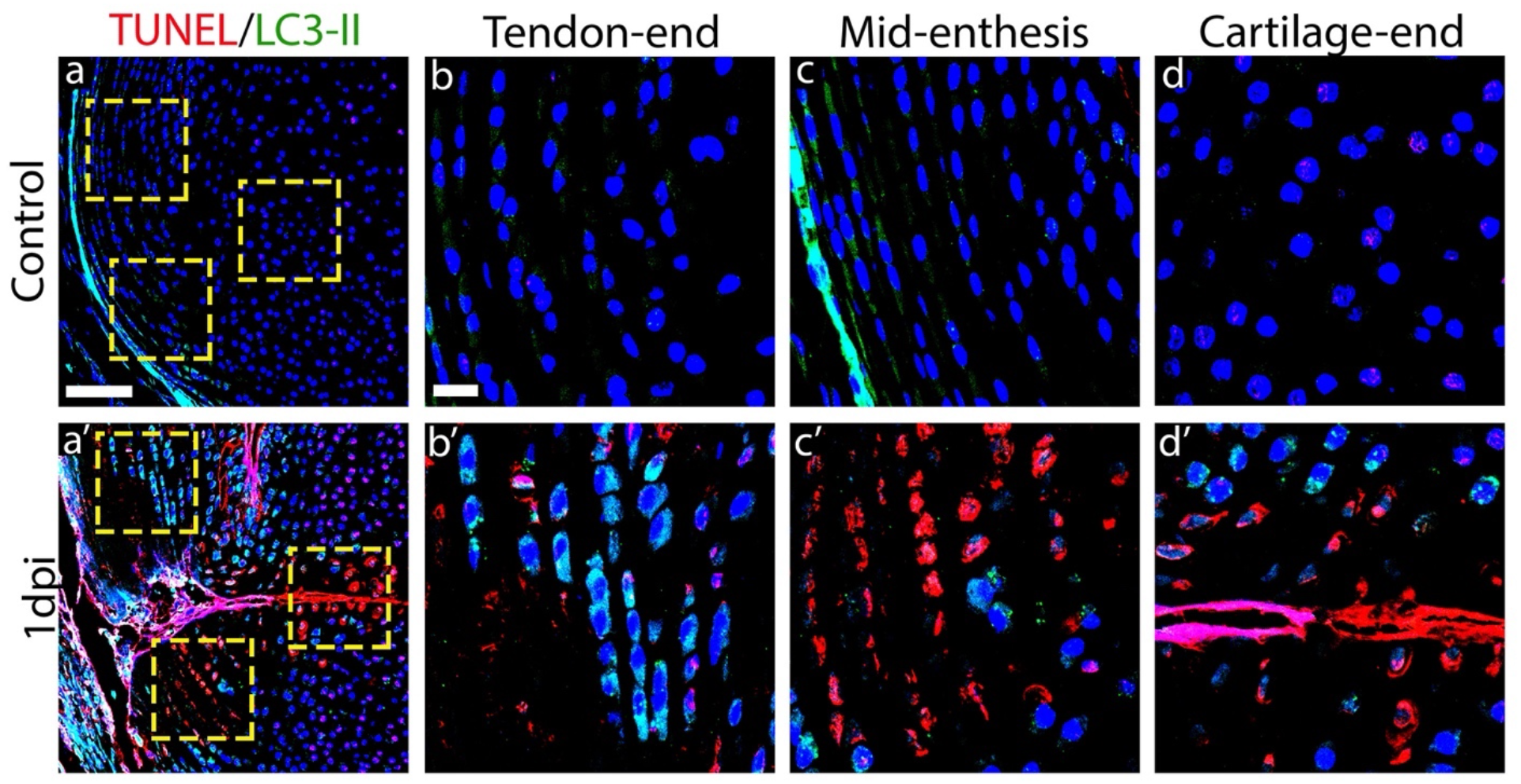
Molecular analyses for autophagy and cell death. TUNEL assay and immunohistochemical staining for autophagy marker LC3-II in control (a) and 1-dpi mouse (a’) entheses. Scale bar: 100 μm. Magnifications of boxed areas show autophagic cells (marked in green) at the tendon end (b’) and near the injury site, along with apoptotic cells (marked in red), which were mainly located in mid-enthesis and cartilage end (c’,d’). Scale bar: 20 μm.

Previous studies of injured articular cartilage describe a cell death process known as chondroptosis. This variant of apoptosis is characterized by an increase in amount of Golgi apparatus and ER, autophagic vacuoles, patchy condensations of nuclei and blebbing of cytoplasmic material^17^, TUNEL-positive cells^36^, and activation of caspase-3 and caspase-9^37^. Because our TEM analysis revealed several of these features, we examined the injury site for cell death by applying TUNEL assay to sections of 1-dpi entheses. At the tendon end, we observed a few cells that were TUNEL-positive; however, the signal was dramatically increased in mid-enthesis and on the cartilage end. Next, we performed immunostaining for cleaved (i.e., activated) caspase-3 (Fig S3); however, we could not detect any signal in or around the injury site. Cellular morphology suggested that the death mechanism was similar to chondroptosis; however, the lack of caspase-3 cleavage suggests that these cells activate a different cell death program.

Together, these results suggest that the different compartments of the injured enthesis undergo both autophagy and cell death.

### Healing of neonatal enthesis via formation of hypocellular domains flanking an ECM plug restored the functional properties of the joint

To determine whether enthesis healing resulted in return of joint function, we next analyzed gait and stride in needle-punched and sham-operated animals at 14, 28 and 56 days post-injury using the CatWalk system. As depicted in Figure 6, at 14 dpi, the injury affected motility, as indicated by a significant reduction in maximum contact area ratio (<1; and compared with sham mice), indicating avoidance from using the paw of the injured leg. This difference was reduced at 28 dpi and by 56 dpi, the injured and sham mice displayed similar results, indicating complete restoration of gait. These results suggest that the healing process effectively restored the function of the injured enthesis.

**Figure 6.**
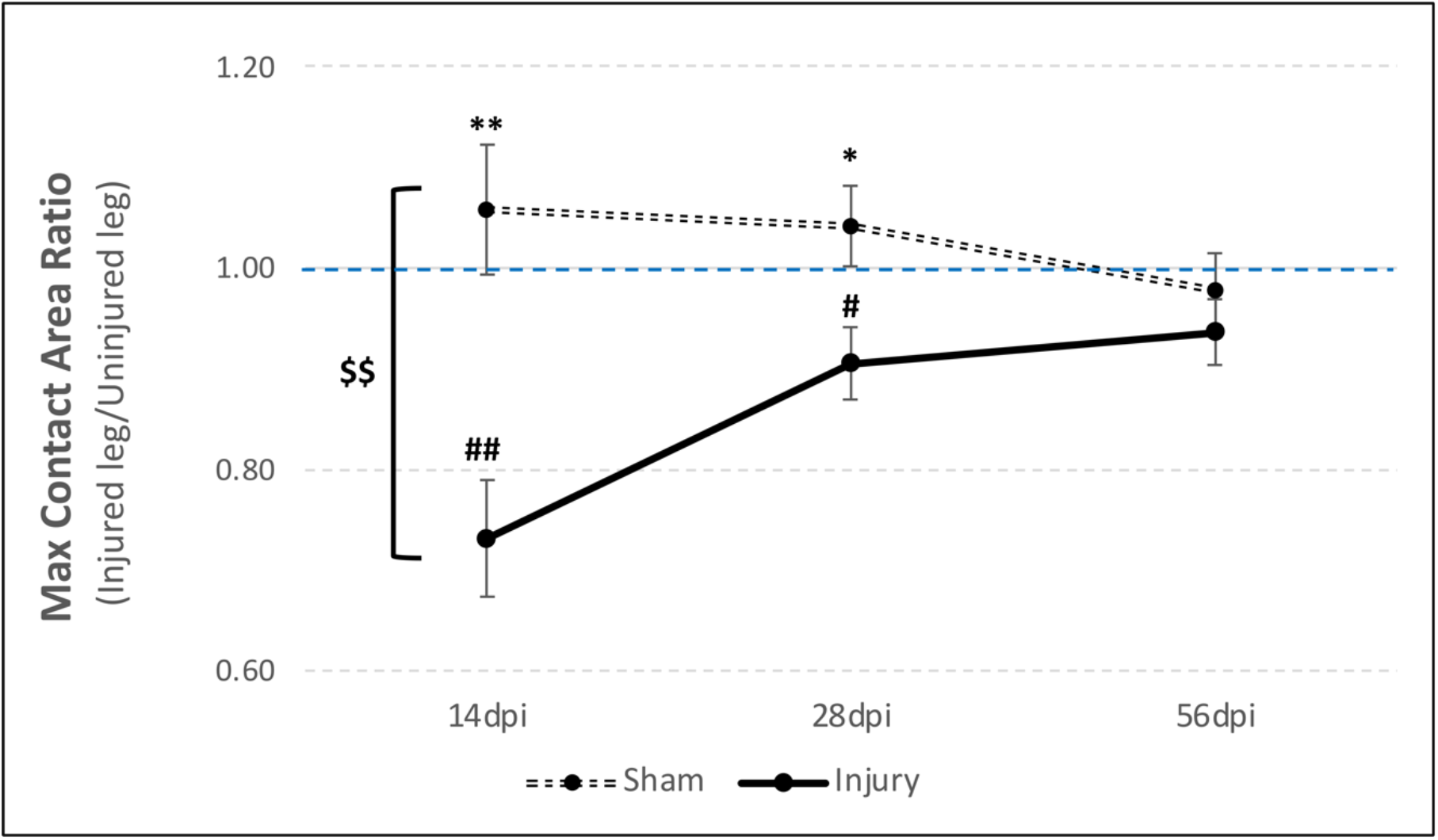
Enthesis healing restores its functional properties. Statistical analysis of CatWalk gait analysis of injury and sham animals at 14, 28- and 56 days after injury (n=10). Data are presented as mean ± SEM. $$, p<0.01 between groups (ANOVA); */**, p<0.05/0.01, respectively, between sham animals (one–sample *t*-test); #/##, p<0.05/0.01, respectively, between injury animals (independent–samples *t*-test).

## DISCUSSION

In this work, we induced partial injuries to Achilles entheses of neonatal mice to uncover the ensuing healing sequence in the different regions of this complex organ. We found that restoration of enthesis function was correlated with an extensive local secretion of ECM, which formed a plug that sealed the lesion, concomitant with cell loss that resulted in an acellular scar tissue flanking the lesion (Fig. 7A). Interestingly, the cellular response to the injury varied along the enthesis. Whereas at the tendon end, most cells underwent ER stress and autophagy, few cells in the cartilage end underwent these processes. Conversely, cell death was more prominent at the cartilage end (Fig 7B). These results suggest a differential cellular response along the healing enthesis.

**Figure 7.**
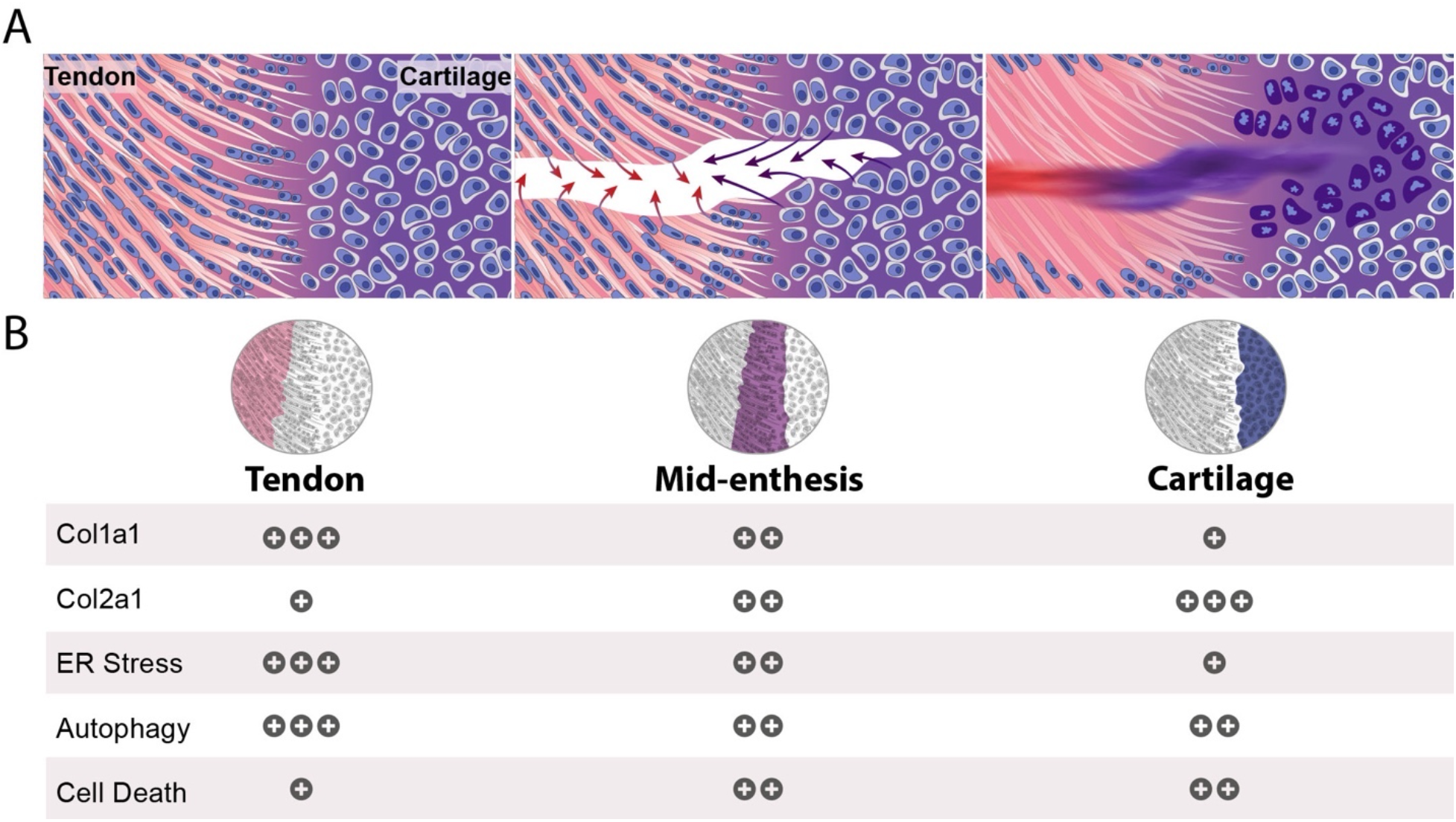
Variations in cellular response between the different regions of the injured neonatal enthesis during the formation of an acellular scar. **A** Illustrations of the neonatal enthesis before (left) and after a needle-punch injury. Shortly after the injury, resident cells adjacent to the injury site secrete ECM corresponding to their cellular identity (middle). This process terminates in the formation of an acellular scar (right, healed enthesis). Red arrows mark Col1a1 and purple arrows mark Col2a1. **B** Spatial distribution of ECM components and molecular markers following injury. Relative levels of expression are indicated by the number of plus signs.

In adult tissues, wound healing typically involves infiltration of phagocytes and fibroblasts into the injury site, where they secrete pro-inflammatory cytokines, chemokines, and growth factors to promote phagocytosis, angiogenesis, cell proliferation, and deposition of collagen, predominantly COL3A1^38^. Usually, this process terminates by formation of a hypercellular fibrovascular scar ^1,2,39,40^. Our findings in the injured neonatal enthesis deviate from the common model in several ways. First, there was no recruitment of blood vessels, immune cells, or fibroblasts to the injury site. Second, the composition of the scar tissue differed from the typical adult fibrovascular scar, as it was hypocellular and its ECM content changed from COL2A1 at the cartilage side to COL1A1 at the tendon side. In the transition area, a mixture of both collagen types was found. Third, the secretory cells that formed the scar tissue were resident cells of the enthesis. Fourth, the process involved ER stress, autophagy, and cell death of the resident secretory cells.

The identity of signals that induce the healing process and the extensive ECM secretion is still an open question. These signals may be molecular, mechanical, or a combination of the two. Putative molecular signals that may drive the process were previously observed in vertebrate and in Drosophila^41–44^. In both cases, the apoptotic cells produced a signal that propagated away and could induce either survival or apoptosis in neighboring cells. However, the enthesis is also a mechanosensitive tissue and, therefore, healing may have also been affected by biophysical forces^45–47^. Moreover, developmental studies have shown that mechanical load is essential for the formation of a proper enthesis^48–50^. It is possible that because of the injury, the loss of tissue integrity alternates the mechanical signals that the enthesis cells sense, leading to their activation. Another interesting characteristic of the healing process observed in the current study is the propagation of the injury signal several rows of cells away from the margins of the injury site.

Whether this propagation was mediated by the original signals that initiated the healing process or by another mechanism requires further investigation.

We showed that during the healing process, resident cells activate ER stress, autophagy, and cell death programs in a region-specific manner. ER stress is commonly associated with extensive ECM production, as the protein load on the ER exceeds its folding capacity^51^. Although this stress response is a coping mechanism, prolonged ER stress can disturb cellular homeostasis and cause cell death^52,53^. In addition, in response to ER stress, autophagy may be triggered to restore homeostasis and to provide an alternative source of intracellular building blocks and energy to the cell^51^. While in some cases autophagy following ER stress has a pro-survival effect, in other cases it promotes self-consumption and cell death^54–56^. Programmed cell death pathways, which are identified by morphological and molecular markers, can be triggered by different stimuli^57^. In the current study, although the dying cells in the cartilage end displayed features of chondroptosis, other characteristics were missing. Therefore, the cell death pathway that we observed may be novel. Different responses were seen on the tendon end compared to the cartilage end of the healing enthesis. The epistatic relations between these cellular phenotypes and their role in the healing process are unknown. Nevertheless, their distribution suggests that the different cell types of the enthesis have different coping strategies in response to injury.

Previous studies have shown that in the adult enthesis, the healing process is characterized by the formation of a hypercellular and vascularized scar^28,58,59^. It is plausible that many of the differences between our observations and previous ones could be driven by age differences. Yet, other factors might contribute to the different outcomes as well. We used a needle punch to injure the enthesis and took great care not to reach the marrow space. In contrast, previous studies have included partial or full detachment of the tendon, reattachment of the tendon to bone, or punch defects that entered the marrow space ^29,60^. Furthermore, injury to deeper structures such as the supraspinatus tendon enthesis requires dissection of surrounding muscles, which can trigger bleeding and infiltration of cells and factors derived from the vasculature^29^. These extrinsic factors may complicate the interpretation of healing mechanisms. In the current neonatal Achilles enthesis injury model, we failed to observe any signs of inflammation at the injury site, a result likely resulting from the subcutaneous position of the enthesis, which lacks overlying musculature, and the poor vascularization^27,46^. These local conditions may limit the ability of chemotaxis and infiltration of immune cells and fibroblasts into the injury site. The effect of local conditions on the healing process was previously observed in a comparison between two different anatomic location of canine flexor tendon, namely the poorly vascularized intrasynovial part, and the well-vascularized extrasynovial part. Whereas injury to the extrasynovial part led to angiogenesis, inflammation and repair, the healing intrasynovial part showed minimal vascularization and muted inflammation, resulting in poor repair^8^.

Finally, although we did not perform mechanical testing of the healed enthesis, gait analysis demonstrated that the mice regained normal motility by two months post injury. This suggests that despite the unusual nature of the healing process and the resulting acellular scar, the healing process produced a mechanically competent tissue. Nonetheless, the injury was created prior to enthesis mineralization, so we cannot exclude the possibility that enthesis mineralization during postnatal maturation contributed to the restoration of mechanical integrity.

In conclusion, our findings provide insight into the healing of the injured neonatal enthesis and present a novel healing mechanism. The key feature of this healing process is the formation of a hypocellular scar by resident cells. The finding of different cellular responses along the length of the enthesis reveals a new level of complexity of this transitional tissue. Recently, the neonatal mouse has emerged as a model for improved healing capacities in diverse mammalian tissues as compared to adults, in which injury often fails to heal or terminates with a fibrotic scar^10,30,61–63^. Identifying the mechanisms that regulate this new healing process may therefore lead to the development of new approaches to the treatment of adult tendon enthesis injuries.

## Authors contribution

**R.C.V** Conceptualization, Methodology, Investigation, Formal Analysis, Visualization and Writing Original Draft. **N.F** Conceptualization, Methodology, Investigation. **E.A**, **E.K** and **S.K** Investigation. **S.R** Methodology. **L.H.Y** Formal Analysis. **N.D** and **S.L.Z** Investigation. **M.T** Investigation and Formal Analysis **S.T** Conceptualization, Writing Original Draft and **E.Z** Conceptualization, Methodology, Supervision, and Writing Original Draft.

## Acknowledgments

We thank Nitzan Konstantin for expert editorial assistance, Prof. Zvulun Elazar from the Department of Biomolecular Sciences, Weizmann Institute of Science, who kindly provided us with GFP-LC3#53 mice, Dr. Ron Rotkoff from the Bioinformatics Unit, Weizmann Institute of Science, for his help with statistical analysis, Tali Wiesel and Tal Bigdary from the Graphic Design Department for their help with graphics and Aaron Vinestock for his assistance in smFISH quantification. Special thanks to all members of the Zelzer Laboratory for encouragement and advice. This study was supported by grants from the National Institute of Health (grant No. R01AR055580) (to S.T. and E.Z.), Israel Science Foundation (ISF, grant No. 1462/20), Minerva Foundation (grant No. 713533), the David and Fela Shapell Family Center for Genetic Disorders, and by The Estate of Mr. and Mrs. van Adelsbergen (to E.Z.).

## METHODS

### Mice

All experiments involving mice were approved by the Institutional Animal Care and Use Committee (IACUC) of the Weizmann Institute. Histology was performed on C57BL6 wild-type mice. GFP-LC3#53 mice were kindly provided by Professor Zvulun Elazar, Weizmann Institute, Israel.

### Achilles enthesis injury model

P7 neonatal mice were anesthetized by lidocaine (0.03 mg, IP). A small incision was made through the skin to expose the Achilles enthesis and was needle-punched using a 32-gauge needle using a sterile approach. The skin was then sutured with nylon 5/0 monofilament. The left limb was used as a control. After injury, the animals returned to full cage activity. Male and female mice were distributed evenly.

### Histological analysis, TUNEL, and Von Kossa staining

For histology, postnatal mice were harvested at various ages, dissected, and fixed in 4% paraformaldehyde (PFA)/ PBS at 4°C overnight. After fixation, tissues were dehydrated to 70% EtOH and embedded in paraffin. Pup and adult tissues were decalcified using 0.5 M EDTA (pH 7.4) prior to dehydration. The embedded tissues were cut to generate 7-μm-thick sections and mounted onto slides.

Hematoxylin and eosin (H&E) and Safranin O stainings were performed following standard protocols. TUNEL assay was performed using In Situ Cell Death Detection Kit (Roche) according to the manufacturer’s protocol.

For Von Kossa and toluidine blue (pH 6.0) staining, fixed calcified tissues were embedded in OCT. 10-μm-thick cryosections were prepared using the Kawamoto film method (Kawamoto and Kawamoto, 2014), and staining was performed using the standard protocols^64^. In short, postnatal mice were harvested at 5 months post-injury, dissected, and fixed in 4% paraformaldehyde (PFA)/ PBS at 4°C overnight. After fixation, tissues were transferred to 30% sucrose overnight, then embedded in OCT and sectioned by cryostat at a thickness of 10 μm. Slides were incubated in 1% silver nitrate solution for 2.5 minutes in UV table, then rinsed 3 times in distilled water. Unreacted silver was removed by 5 minutes incubation in 5% sodium thiosulfate, then rinsed in water. Then, sections were counterstained with toluidine blue. Lastly, slides were mounted with Entellan (Sigma-Aldrich, 1079600500).

### Immunofluorescence

For immunohistochemistry on paraffin sections, animals were harvested at various ages, dissected, and fixed in 4% PFA/PBS at 4°C overnight. After fixation, tissues were decalcified using 0.5 M EDTA (pH 7.4), washed thoroughly with water, dehydrated to 70% EtOH and embedded in paraffin. The embedded tissues were cut to generate 7-μm-thick sections and mounted onto slides. Antigen retrieval for anti-collagen types I, II and III antibodies was performed using 1.8 μg proteinase K (Sigma-Aldrich, P9290) in 200 mL PBS for 10 minutes. Antigen retrieval for anti-GFP, F4/80 and cleaved caspase-3 antibodies was performed in 10 mM sodium citrate buffer (pH 6.0) cooked in 80°C for 15 min in hot tub. Then, sections were washed twice in PBS and endogenous peroxidase was quenched using 3% H_2_O_2_ in PBS. Non-specific binding was blocked using 7% horse serum and 1% BSA dissolved in PBST for 1 hour. Then, sections were incubated with either rabbit anti-collagen I antibody (1:100, # NB600-408, Novus biologicals), mouse anti collagen II antibody (1:50, II-II6B3, the Developmental Studies Hybridoma Bank), rabbit anti-collagen III (1:100, ab7778,Abcam), goat anti-GFP (biotin) antibody (1:100, ab6658, Abcam), rabbit anti-cleaved caspase-3 (Asp175) antibody (1:200, #9664s, Cell Signaling) or rat anti-F4/80 antibody (1:50, ab6640, Abcam) overnight at room temperature. The next day, sections were washed twice in PBST and incubated with biotin anti-rabbit (1:100 Jackson ImmunoResearch), biotin anti-rat, (1:100 Jackson ImmunoResearch), for 1 hour in room temperature. Then, after two washes of PBST, slides were incubated with streptavidin-Cy2 or streptavidin-Cy3 (1:100, Jackson ImmunoResearch), Cy3-conjugated donkey anti-mouse (1:100, Jackson ImmunoResearch). Occasionally, slides were counterstained using DAPI. Then, slides were mounted with Shandon Immu-mount (#9990402, Thermo-Scientific).

For immunohistochemistry on cryosections, animals were harvested at various ages, dissected, and fixed in 4% PFA/PBS at 4°C overnight. Then, tissues were decalcified using 0.5 M EDTA (pH 7.4) and transferred to 30% sucrose overnight, embedded in OCT and sectioned by cryostat at a thickness of 10 μm. Cryosections were dried and post-fixed for 20 min in acetone at −20°C. Then, sections were permeabilized with 0.2% Triton/PBS. To block non-specific binding of immunoglobulin, sections were incubated with 7% goat serum in PBS. Cryosections were then incubated overnight at 4°C with primary antibody rat anti-mouse CD31 (BD PharMingen, PMG550274; 1:50). The next day, sections were washed in PBS and incubated with biotin anti-rabbit (1:100 Jackson ImmunoResearch). Then, slides were incubated with streptavidin-Cy3 (1:100, Jackson ImmunoResearch) and Cy3-conjugated donkey anti-rabbit (1:100, Jackson ImmunoResearch). Occasionally, slides were counterstained using DAPI. Then, slides were mounted with Shandon Immu-mount (#9990402, Thermo-Scientific).

### Single-molecule fluorescent in situ hybridization

Single-molecule FISH was performed using HCR V3.0, as previously described by Choi et al.^65^ with slight modifications. The probes for *BiP* and *Chop* were designed and ordered from Molecular Instruments. mRNA accession numbers are shown in Table 1. Briefly, tissue was fixed for 3 hours following sacrifice using 4% PFA/PBS freshly prepared with DEPC water. Then, solution was changed to 4% PFA/PBS/ 30% sucrose and was incubated shaking overnight. The following day, the tissue was embedded in OCT and kept at −80°C until use. On the morning of the experiment, the tissue blocks were cut to produce 10-μm thick sections using a cryostat, and kept in the cryo chamber at −30°C until the beginning of the experiment. Then, tissue sections were warmed to room temperature and dried in a chemical hood for 7 min, followed by incubation in 70% EtOH/DEPC at 4°C for 1 hour. Then, sections were washed once in PBS and fixed in 4% RNase-free PFA for 7 min, washed with RNAse-free PBS, and permeabilized in 10 μg/ml PK/PBS for 10 min at room temperature. Section were then washed with PBT (RNAse-free PBS-tween 0.1%) twice, post-fixed with 4% RNase-free PFA for 5 min and washed again with PBT twice for 5 min. Then, sections were washed with acetylation buffer for 10 min, twice with PBT and rinsed in DEPC, as previously described in Shwartz & Zelzer (2014). Next, sections were left to dry for 30 min at room temperature and equilibrated in HCR hybridization buffer (Molecular Instruments) for 10 min at 37°C. Probes were then added to the sections at a final concentration of 0.4-4 nM and hybridized overnight in a humidified chamber at 37°C. The next day, the protocol described by Choi et al.^65^ was applied using home-made wash buffer (50% formamide (Merck Millipore, 75-12-7), 5XSSC (Molecular Biology-P, 001985232300), 9 mM citric acid (pH 6.0) (Sigma Aldrich, 77-92-9), 0.1% Tween20 (Sigma Aldrich, P1279) and 50 μg/ml heparin (Sigma Aldrich, H3393)). The probe sets were amplified with HCR hairpins for 45 min at room temperature in HCR amplification buffer (Molecular Instruments). Fluorescently-conjugated DNA hairpins used in the amplification were ordered from Molecular Instruments. Prior to use, the hairpins were ‘snap cooled’ by heating at 95°C for 90 sec cooling to room temperature for 30 min in the dark. After amplification, the samples were washed in 5X SSCT and stained with DAPI (1:10,000, D9542, Millipore Sigma) diluted in PBS for 5 min and then mounted onto slides using Shandon Immu-mount (#9990402, Thermo-Scientific). Sections were imaged with a confocal LSM 800 microscope (Zeiss) at a resolution of 70 nm (x,y). The brightness of the *in situ* signal was enhanced in FIJI for presentation in the figure. For quantification of transcripts per cell, amplification was performed for 1 hour and sections were imaged with the same microscope at a resolution of 70 nm and 400 nm (x,y,z). Autofluorescence imaging of cells was acquired with excitation at 488 nm laser, and cells were segmented with Cellpose (green), using the cyto algorithm with a diameter size between 75-500 pixels, depending on the area in the enthesis. Images were then further quantified in cellprofiler with custom pipelines.

**Table 1.** List of probes used for *in situ* HCR and their accession numbers.

| Probe name | Accession number | HCR Amplifier | Amplifier color |
| --- | --- | --- | --- |
| BiP | NM_022310.3 | B5 | 546 nm |
| Chop | NM_007837.4 | B3 | 488 nm |

### Transmission electron microscopy

Animals were harvested at 1- and 3-days post-injury, dissected, and fixed with 3% paraformaldehyde, 2% glutaraldehyde in 0.1 M cacodylate buffer containing 5 mM CaCl_2_ (pH 7.4) overnight. The tissue was decalcified using 0.5 M EDTA (pH 7.4) for 96 h. Vibrotome sections (200 micrometer) were prepared (Leica VT1000 S) and tissue was postfixed in 1% osmium tetroxide supplemented with 0.5% potassium hexacyanoferrate trihydrate and potassium dichromate in 0.1 M cacodylate (1 h), stained with 2% uranyl acetate in water (1 h), dehydrated in graded ethanol solutions and embedded in Agar 100 epoxy resin (Agar scientific Ltd., Stansted, UK). Ultrathin sections (70-90 nm) were obtained with a Leica EMUC7 ultramicrotome and transferred to 200-mesh copper TEM grids (SPI). Grids were stained with lead citrate and examined with a FEI Tecnai SPIRIT (FEI, Eidhoven, Netherlands) TEM operated at 120 kV and equipped with a Gatan OneView camera.

### CatWalk gait analysis

Gait and stride were assessed using the CatWalk XT 10.6 automated gait analysis system (Noldus Information Technology, Wageningen, The Netherlands) at 14, 28 and 56 days post-injury. Mice were subjected to at least 5 runs in each assessment session. Following the identification and labeling of each footprint, gait data were generated. The collected data included maximum contact area, which is the area of the foot touching the surface during the time of maximum contact. To quantify functional changes, we divided the maximum contact area of the manipulated hindlimb (right) by that of the intact hindlimb (left). This ratio index was used for quantification and statistical analyses.

### Quantitative real-time (qRT-) PCR

Total RNA was purified from Achilles enthesis of 1-dpi pups or from tissue culture cells using the RNeasy Kit (Qiagen). Reverse transcription was performed with High-Capacity cDNA Reverse Transcription Kit (Applied Biosystems) according to the manufacturer’s protocol. For control, mouse embryonic fibroblasts were treated with Brefeldin A (B7651-5MG, Sigma Aldrich). qRT-PCR was performed using Fast SYBR Green master mix (Applied Biosystems) on the StepOnePlus machine (Applied Biosystems). Values were calculated using the StepOne software version 2.2, according to the relative standard curve method. Data were normalized to 18S rRNA. Primer sequences are given in Table 2.

**Table 2.**
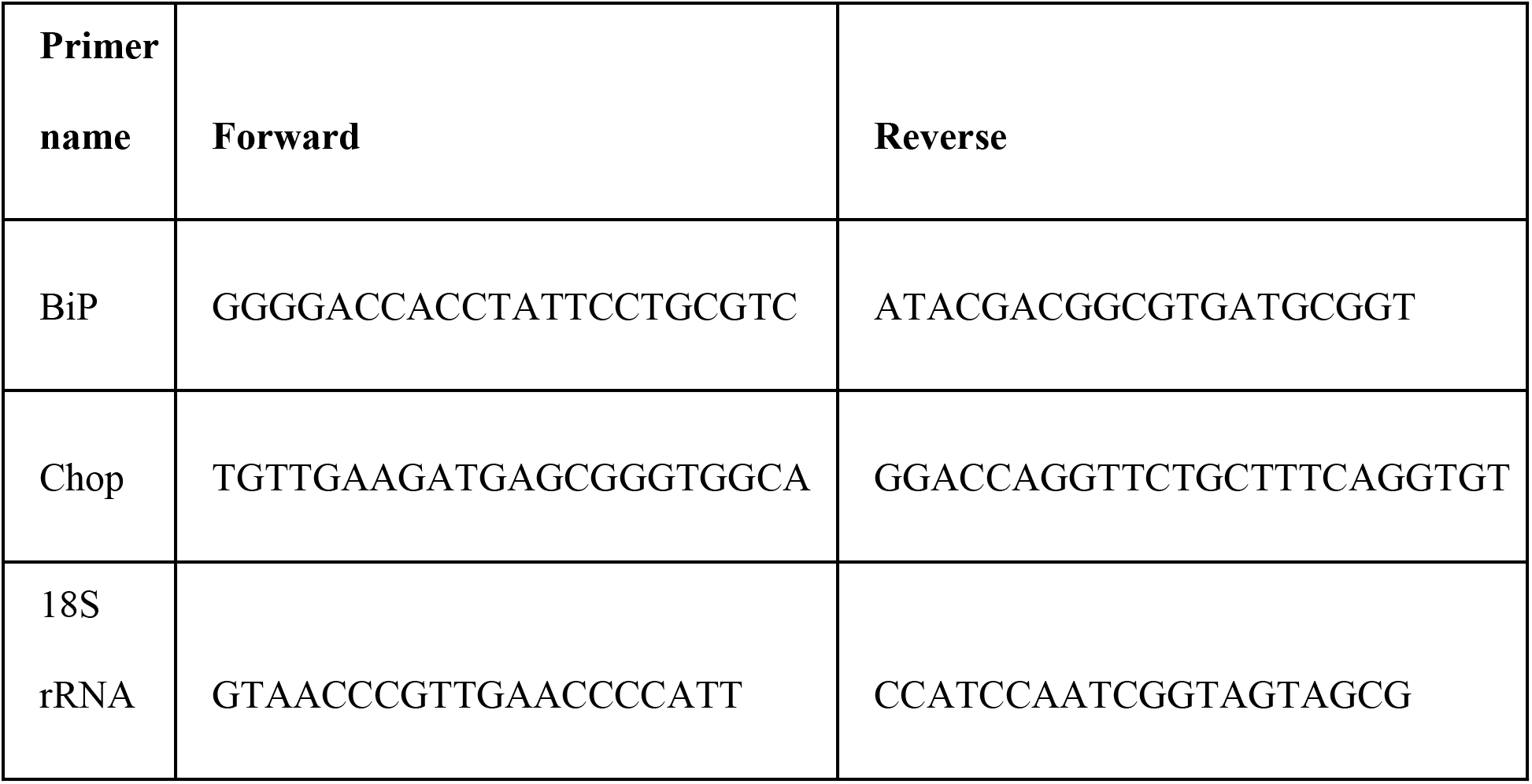
List of primers used for the qRT-PCR and their sequences.

### Statistical analysis

Statistical analyses of qRT-PCR results was performed with Excel using paired two-tailed Student’s *t*-test. CatWalk gait analyses were performed using SPSS two-way ANOVA and One-sample *t*-test. Quantification of single-molecule FISH HCR results was performed using linear mixed model. The data are presented as mean±SD. All statistical details, including n values, are given in the figures and figure legends.

## Supplementary Figures

**Supplementary Figure 1.**
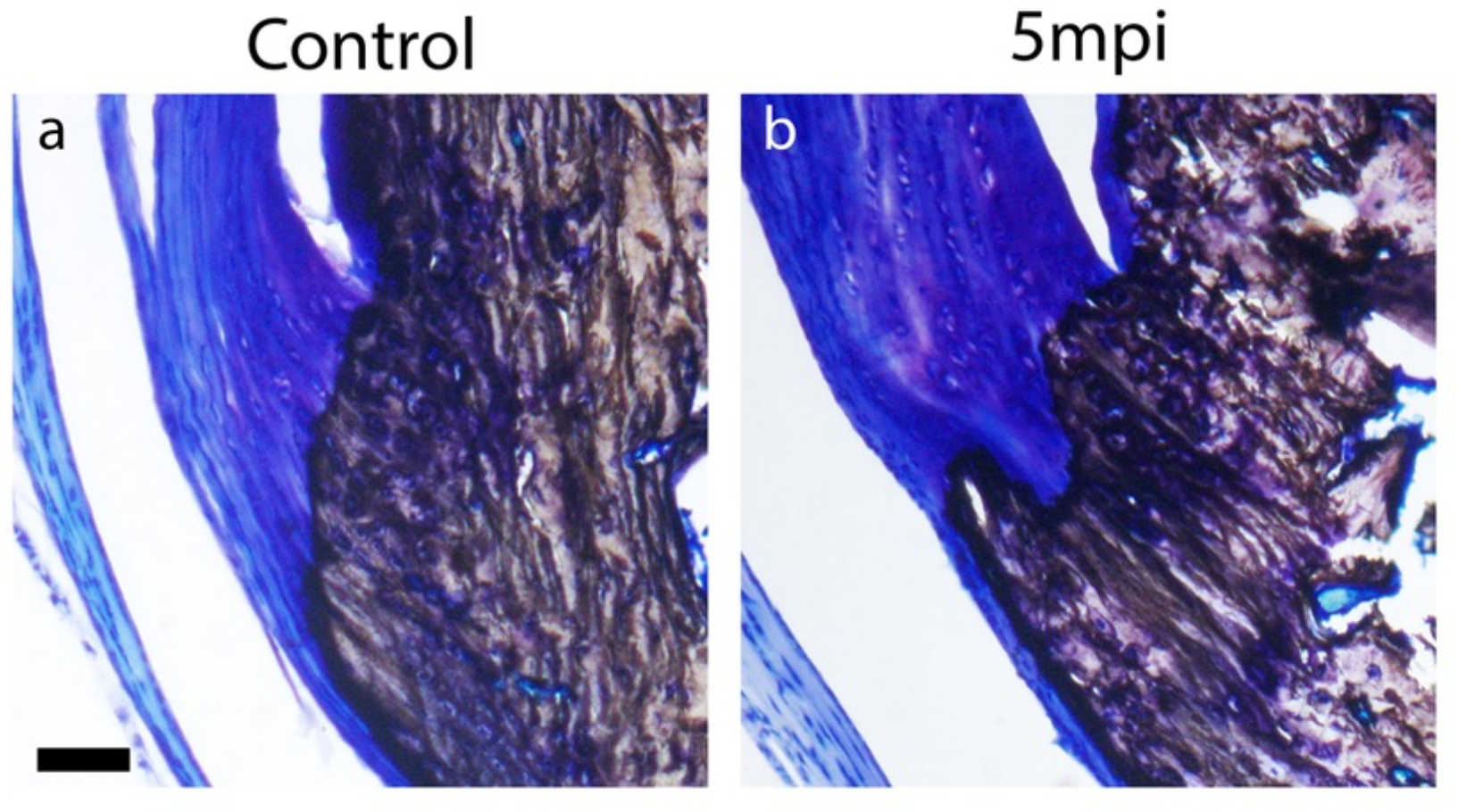
Injured enthesis exhibits a disrupted tidemark as the ECM plug does not undergo mineralization. Von Kossa staining of sagittal sections from control uninjured leg (a) and injured entheses at 5 months post-injury (mpi) (b). Scale bar: 50 μm.

**Supplementary Figure 2.**
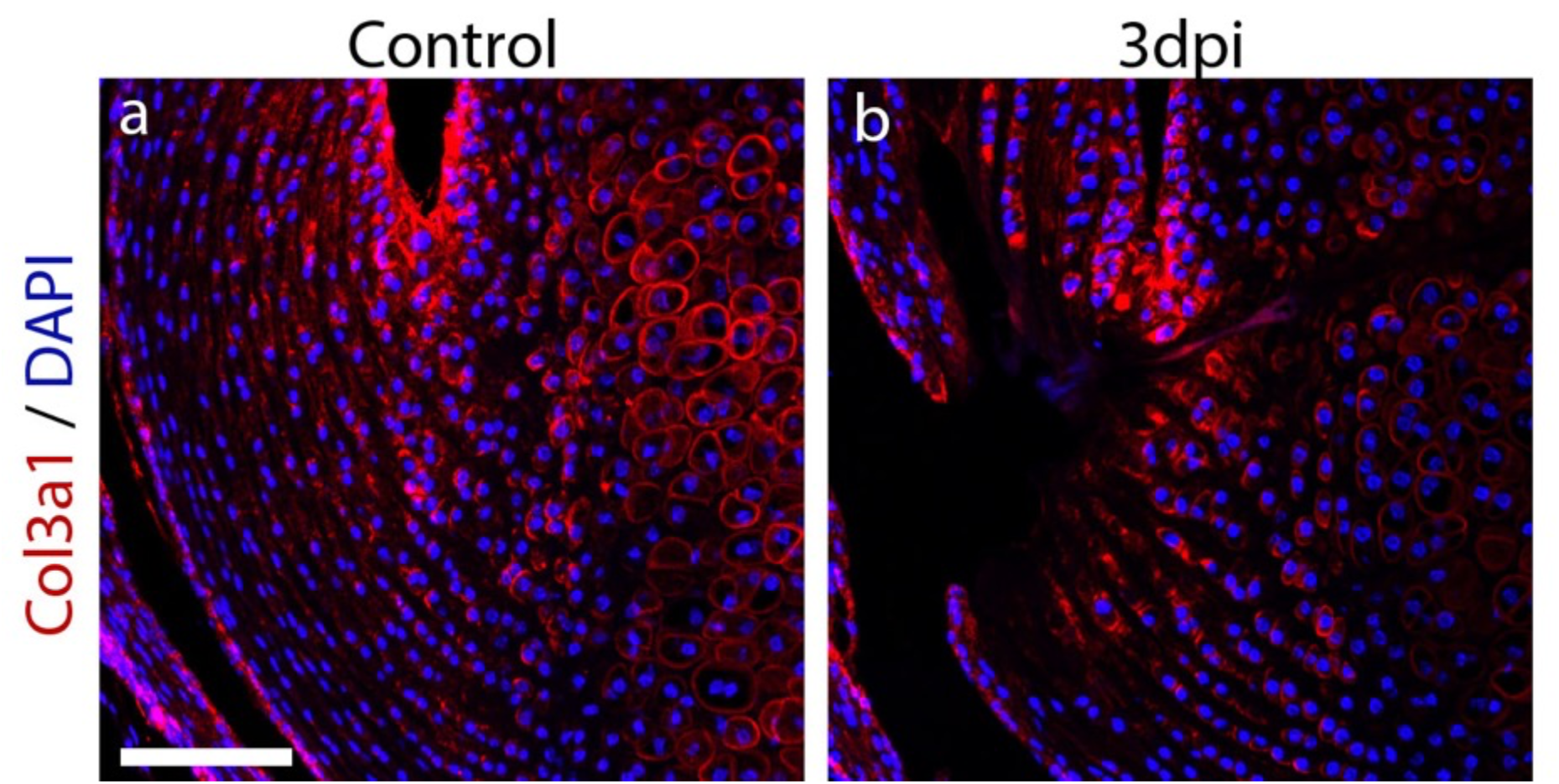
Collagen type III is absent from the ECM plug in the injured enthesis. Immunohistochemistry staining for COL3A1 of control (a) and 3-dpi (b) entheses shows no change in collagen composition in the ECM plug. Scale bar: 100 μm.

**Supplementary Figure 3.**
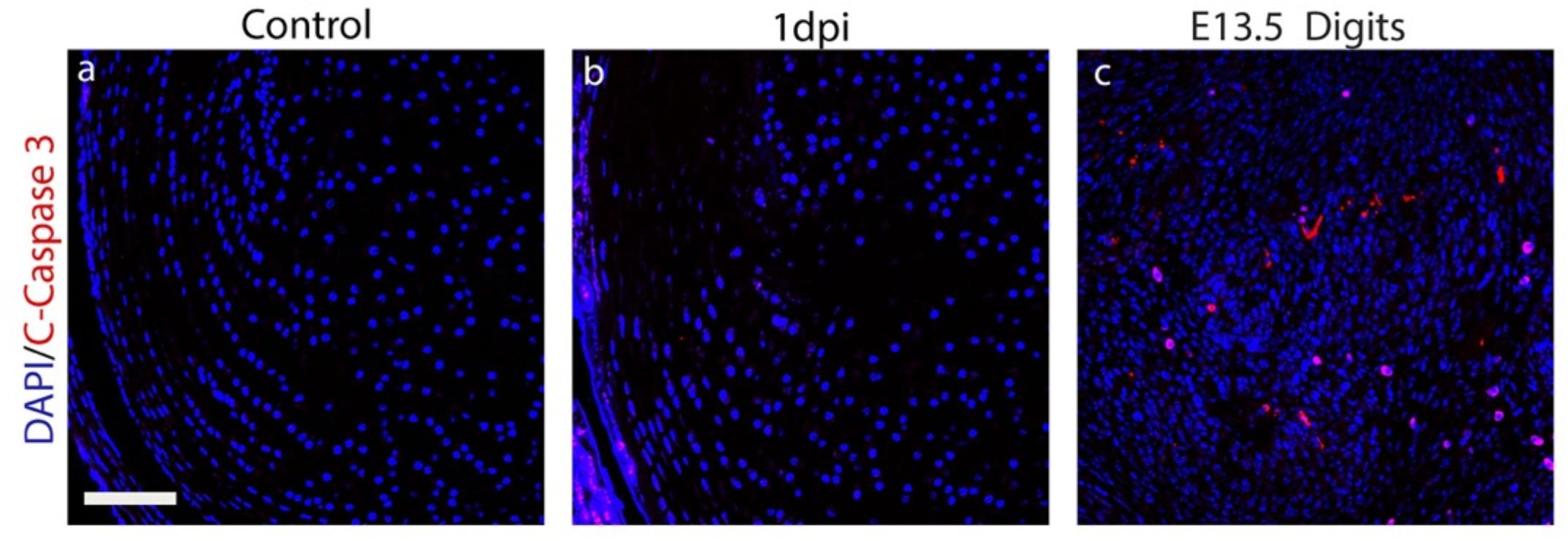
Cleaved caspase-3 is absent from the ECM plug in the injured enthesis. Immunohistochemistry staining of control (a) and 1-dpi entheses (b) shows no presence of cleaved caspase-3 in proximity to the injured enthesis. E13.5 digits section was used as a positive control (c). Scale bar: 100 μm.

**Supplementary Figure 4.**
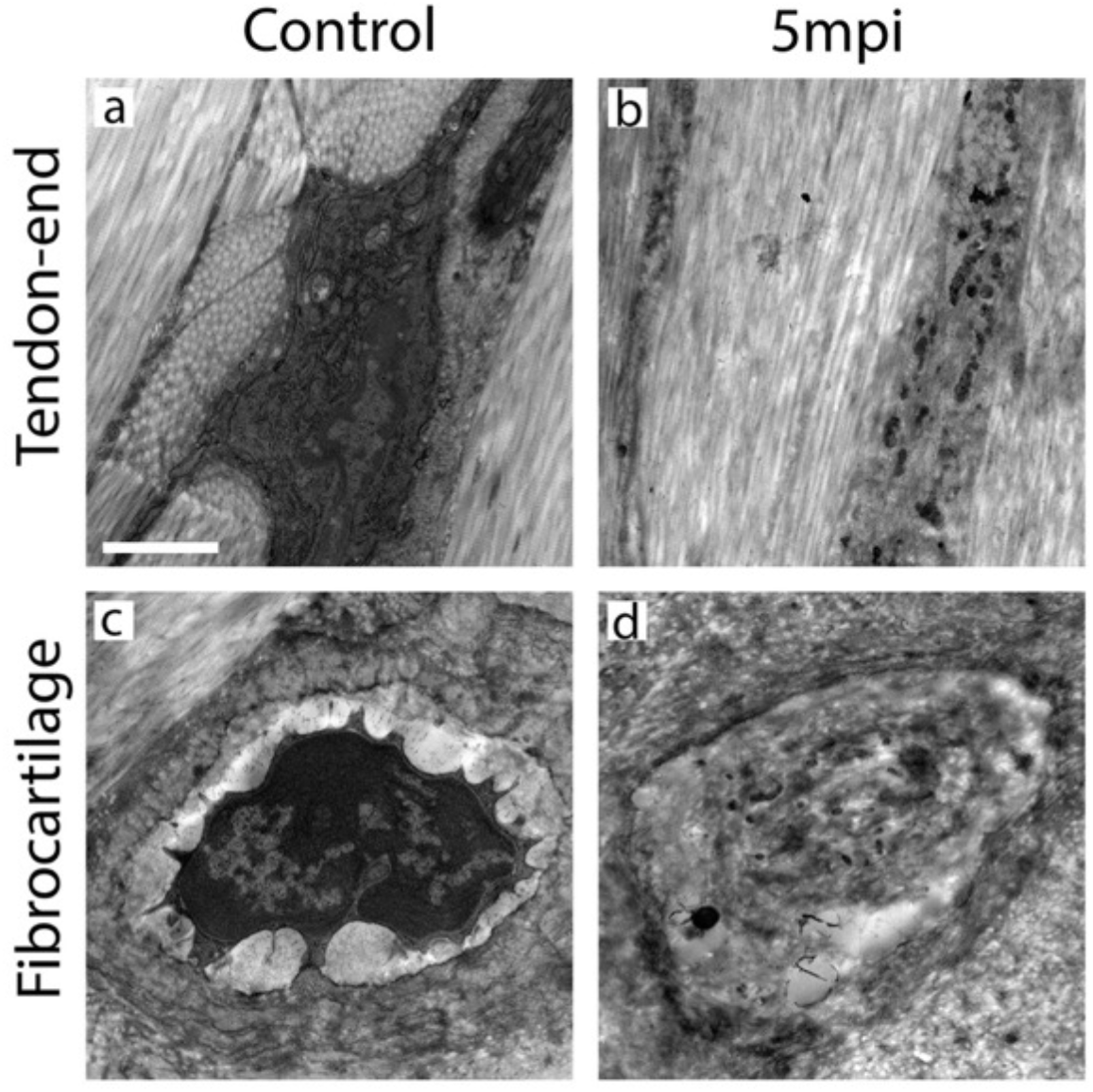
TEM analysis reveals a long-term impairment in cell morphology in the hypocellular scar of the injured enthesis. TEM images show cellular morphology of control and 5 months post-injury entheses at the tendon end (a-b) and cartilage end (c-d). Scale bar: 2 μm.

